# Divergence in gut microbial communities mirrors a social group fission event in a black-and-white colobus monkey (*Colobus vellerosus*)

**DOI:** 10.1101/511618

**Authors:** Claire K. Goodfellow, Tabor Whitney, Diana M. Christie, Pascale Sicotte, Eva C. Wikberg, Nelson Ting

**Author notes:** Wikberg and Ting should be considered co-senior/co-corresponding authors.

## Abstract

Host behavior and social factors have increasingly been implicated in structuring the composition of gut microbial communities. In social animals, distinct microbial communities characterize different social groups across a variety of taxa, although little longitudinal research has been conducted that demonstrates how this divergence occurs. Our study addresses this question by characterizing the gut microbial composition of an African Old World monkey, the black-and-white colobus (*Colobus vellerosus*), prior to and after a social group fission event. Gut microbial taxonomic composition of these monkeys was profiled using the V-4 hypervariable region of the bacterial 16s rRNA gene, and pairwise-relatedness values were calculated for all individuals using 17 STR loci and partial pedigree information. The two social groups in this study were found to harbor distinct microbial signatures after the fission event from which they emerged, while these communities were not divergent in the same individuals prior to this event. Three genera were found to differ in abundance between the two new social groups: *Parabacteroides*, *Coprococcus*, and *Porphyromonadaceae*. Additionally, although this fission happened partially along lines of relatedness, relatedness did not structure the differences that we found. Taken together, this study suggests that distinct gut microbial profiles can emerge in social groups in less than one year and recommends further work into more finely mapping the timescales, causes, and potentially adaptive effects of this recurring trend toward distinct group microbial signatures.

**Research highlights:**

- Distinct gut microbial profiles emerge in two social groups of *C. vellerosus* less than nine months after a fission event.
- Three genera differ in abundance between the two new social groups.
- Relatedness does not structure differences in microbial composition between the groups.

## INTRODUCTION

The mammalian gut harbors a dynamic microbial community which contributes to host physiology, metabolism, and defense (Huttenhower *et al*., 2012; Cho & Blaser, 2012; Barbachano *et al.,* 2017). This community both shapes host phenotypes and is shaped by host characteristics that can include phylogeny, genetic variation, environment, and spatial distribution (Wu *et al*., 2011; Leamy *et al*., 2014; Barelli *et al*., 2015; Amato, 2013; Amato *et al*., 2016). Moreover, behavior and social context can contribute to gut microbiome composition, and diet choice, habitat use, mate choice, and social networks have all been shown to modulate gut diversity (Archie *et al*., 2015; Ezenwa *et al*., 2012). In fact, the social transmission of beneficial microbes has been cited as one of the benefits associated with group living (Lombardo, 2008), and it has been suggested that more similar gut communities between hosts may confer similar “ecosystem services” to their hosts (Costello *et al*., 2012). Given that conspecifics in the same social group likely encounter highly comparable ecological challenges, the social transmission and ultimate convergence of a group’s gut microbiota into the consortia that provides the most ideal ecosystem services for that particular group’s set of demands could prove to be of evolutionary benefit.

Distinguishing the complex and intertwined forces that shape this dynamic community, however, is difficult. Studies of wild populations can help to address this difficulty, providing insight into the forces at play in natural communities as well as how they change over time (Amato, 2013). In particular, studies of wild primates and other highly social animals allow us to answer important questions about how social forces shape these changes in some of our own closest living relatives. For example, in a number of wild primate populations, more closely associated individuals have more homogenous gut microbiome compositions (Perofsky *et al*., 2017; Moeller *et al*., 2016; Amato *et al.,* 2017) and distinct gut microbiota characterize different social groups in the same population (Tung *et al.,* 2015; McCord *et al*., 2014; Bennett *et al*., 2016; Springer *et al*. 2017; Degnan *et al.,* 2012). Taken together, these findings highlight the importance of social context in gut microbiome composition. Furthermore, immigrant males which have resided in a new social group for a longer period of time have more similar gut microbiota to the resident males of that group, suggesting that the convergence of group member microbiota may occur over a span of months to years (Grieneisen *et al*., 2017). Because dietary shifts typically alter the composition of microbial communities over a shorter time scale of days to weeks, this finding suggests that distinct group communities are not solely the result of changes in diet (Bonte *et al*., 2012; Williams *et al*., 2012; Turnbaugh *et al*., 2009). However, further studies are needed to more thoroughly examine the time scales over which these convergences occur in natural communities.

Group fission events provide an ideal natural system for interrogating such questions. These are a means of group proliferation in social animals that occur when the costs of living in a certain group have grown to outweigh the benefits (Sueur & Maire, 2014). When this happens, one or more social groups will break off from the original group, oftentimes splitting along lines of relatedness (Widdig *et al*., 2006; Snyder-Mackler *et al*., 2014). This type of event provides us the opportunity for unique insight into the physiological and behavioral changes in individuals following such an event as well as the time scales over which they occur. For example, fission events and variations in group size have enabled insight into the effects of social context on grooming networks, fertility and cortisol levels in primates (Henzi *et al*., 1997; Dunbar, 2018; Markham *et al*., 2015). Here, we report on the gut microbiome compositions of a group of ursine colobus or white-thighed black-and-white colobus (*C. vellerosus*) before, and less than a year after, a fission event. The aim of this study was to examine the plasticity of the gut microbiome shortly following a fission event as a way of gaining insight into the time scale over which microbiomes diverge into the distinct microbiomes that have been shown to characterize different social groups.

## METHODS

### Study system

The Boabeng-Fiema Monkey Sanctuary (BFMS) is a 1.92 km^2^ dry semi-deciduous forest (Hall & Swaine, 1981) located in central Ghana (7°43’ N and 1°42’W). Ursine colobus or white-thighed black-and-white colobus (*Colobus vellerosus*) is one of two diurnal primate species that resides at BFMS (Saj *et al*., 2005). This is an arboreal, folivorous monkey (Saj & Sicotte 2007a, 2007b) that lives in uni-male or multi-male multi-female groups of 9-38 animals (Kankam & Sicotte, 2013; Wong & Sicotte, 2006). Dispersal is male-biased in this species (Teichroeb *et al*., 2011), although females do show facultative dispersal (Teichroeb *et al*., 2009; Teichroeb *et al*., 2011; Wikberg *et al*., 2012; Sicotte *et al*., 2017). Female social networks are affected by the presence of infants, kinship, and immigration status, but not by dominance rank (Wikberg *et al*., 2013; Wikberg *et al.,* 2014a; Wikberg *et al*., 2014b; Wikberg *et al*., 2015). Several groups of *C. vellerosus* have been followed systematically since 2000 for behavioral, demographic and ecological data (see for example Teichroeb & Sicotte, 2009; Bădescu *et al*., 2015). Fecal samples were collected on a regular basis from each focal female in our study groups between 2006 and 2009 (see for example Wikberg *et al*., 2015). This study follows the fission of one social group into two daughter groups (named NP and DA) over the course of one year (2006-2007).

### Ethical note

This study was approved by the University of Calgary’s Animal Care Committee, and conducted with permission from the Ghana Wildlife Division and the management committee at BFMS. This research adhered to the American Society of Primatologsts’ Principles for the Ethical Treatment of Non Human Primates.

### Sample collection and STR genotyping

All fecal samples from the study period were collected using masks, fresh gloves, sterile sticks, and sterile tubes to minimize contamination. 1-2 grams of feces were mixed with 6 µL of RNAlater® (Thermo Fisher Scientific, Waltham MA) immediately upon collection and stored at −20°C in the field. After shipment to the Ting lab, samples were again stored at −20°C until DNA extraction. DNA was extracted from two or more samples of each individual using a QIAamp DNA Stool Mini Kit with a slightly modified manufacturer protocol (Wikberg *et al*., 2012), and negative controls were processed with each round of extraction. 17 short tandem repeat (STR) loci were amplified using Qiagen’s multiplex PCR kit with a modified protocol and analyzed on an ABI 3730 DNA analyzer (following Wikberg *et al*., 2012). We determined how many replicates were needed to confirm homozygote genotypes based on real-time PCR DNA quantification (Morin *et al*., 2001). Two replicates were used to confirm heterozygote genotypes.

### Gut microbial profiling

We collected metagenomic data from matched genotyped samples from female members of the original social group collected from June to August, 2006 (n = 12 samples) and from matched genotyped samples from the same female individuals residing in the two daughter groups from July to August, 2007 (NP with n = 6 samples; DA with n = 6 samples). DNA was extracted again as above and quantified using a Qubit dsDNA BR Assay Kit protocol using a Qubit 2.0 Fluorometer (Thermo Fisher Scientific, Waltham MA). Samples containing at least 1.0 ng/µL were chosen for preparation and sequencing of the V-4 hypervariable region of the bacterial 16S ribosomal RNA gene in the Genomics and Cell Characterization Core Facility at the University of Oregon. 200 ng of DNA diluted in 10 µL of H_2_O were PCR amplified using barcoded Illumina 515F and 806R primers. Targets were amplified in reactions of 1 µL DNA, 1.25 µL of 10 µM primer mix, 10.25 µL H2O, and 12.5 µL NEB Q5 hot start 2x Master Mix. The thermal cycling profile was as follows: initial denaturing at 98°C for 0:30, 20-30 cycles of 98°C for 0:10, 61°C for 0:20, and 72°C for 0:20, and a final extension of 72°C for 2:00. PCR products were cleaned using Ampure XP beads (Beckman Coulter, Brea, CA), quantified and normalized. Barcoded amplicons were pooled and pair-end sequenced with 150 base pair reads on a partial medium output run on the Illumina NextSeq platform. Sequences were then demultiplexed and denoised using DADA2 (Bolger *et al*., 2014). Taxonomic units were assigned using the Qiime2 pipeline. An OTU table was generated for samples rarefied to an even sampling depth of 46,040 reads per sample, retaining 1,104,960 sequences for 24 samples. Negative controls were processed at both the extraction and library preparation (PCR) stages, and they were sequenced and carried through the data processing pipelines. No evidence of contamination was found via fluorometry or gel electrophoresis during laboratory work, nor was there any evidence of contaminating sequences in the Illumina reads for the negative controls.

### Data Analyses

Unless otherwise noted, all subsequent statistical analyses were run in R. To test whether average gut microbial composition differed by social group, samples were analyzed using four groups based on social group at the time of sample collection: original group which became NP after the fission, original group which became DA after the fission, DA, and NP. Beta diversity was calculated for the samples as Bray-Curtis dissimilarity using the *phyloseq* package for R (McMurdie & Holmes, 2013). This metric was selected over metrics accounting for evolutionary relatedness as it represents a quantitative measure of community dissimilarity based on relative abundance without adjusting for the phylogenetic proximity of OTUs, thereby allowing insight into structure without accounting for relatedness among microbial taxa. We also ran a PERMANOVA using the *adonis* function in the *vegan* package, with social group (pre-fission DA, pre-fission NP, post-fission DA, post-fission NP), age class (subadult, adult), collection site (mature forest, woodland), and reproductive status of the individual (cycling, non-cycling, pregnant, lactating) as predictors in the model. Linear effect size analysis (LefSe) was run for the groups before and after the fission event at a KW alpha value of 0.01 and an LDA score of 3.0 (Segata *et al*., 2011).

To assess whether the fission occurred along lines of host relatedness, we calculated relatedness following Wikberg *et al*. (2012). A Kruskal-Wallis test was run using metrics of average group relatedness for the three social groups (original group, DA, NP). Finally, we determined whether host relatedness explained differences in gut microbial beta diversity seen between groups. We investigated the correlation between the Bray-Curtis dissimilarity matrix and the host relatedness matrix for each group using Mantel tests, and Spearman rank correlation statistics were computed with 999 permutations.

## RESULTS

In the summer of 2006, one of our study groups (28 individuals) showed elevated levels of female aggression. Subgroups started to range 50 meters apart for periods of time, although the subgroups always convened during the day (Wikberg, unpublished data). *C. vellerosus* typically exhibit a smaller group spread, and 50 meters is used to define a between-group encounter in this species (Sicotte & MacIntosh, 2004). The home range of the original group spanned approximately 0.20 km^2^ through both woodland and mature forest. By May of 2007, this group had fissioned into two daughter groups: NP (10 individuals) and DA (18 individuals) (Table 1). These two groups both remained on the original home range, splitting the range after the fission (Fig. 1). Both daughter groups ranged in subsets of the original range that included both woodland and mature forest, but DA ranged 0.15 km^2^, moreso in woodland areas, while NP’s new range was 0.054 km^2^, moreso in mature forest after the fission event.

**Table 1.**
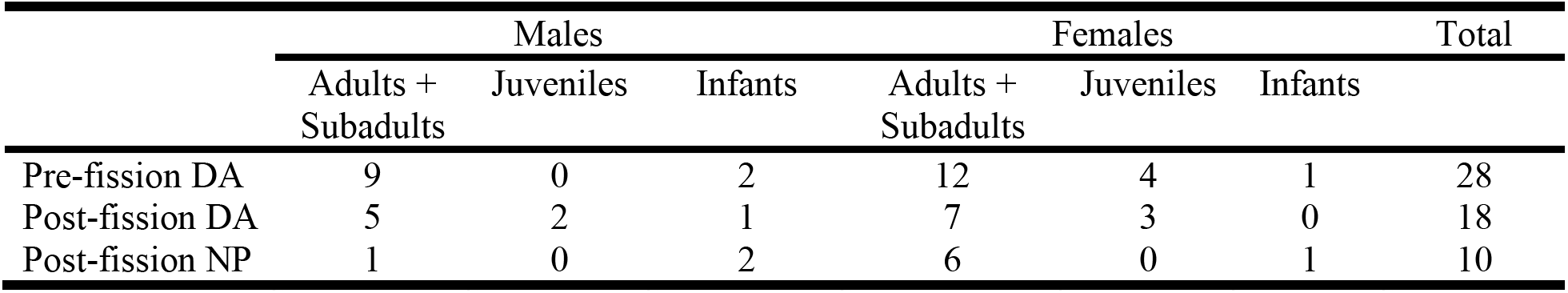
Composition of the different social groups before and after the fission event. Although this study only examined the gut microbial compositions of females, males are included in this table for insight into group composition overall.

**Figure 1.**
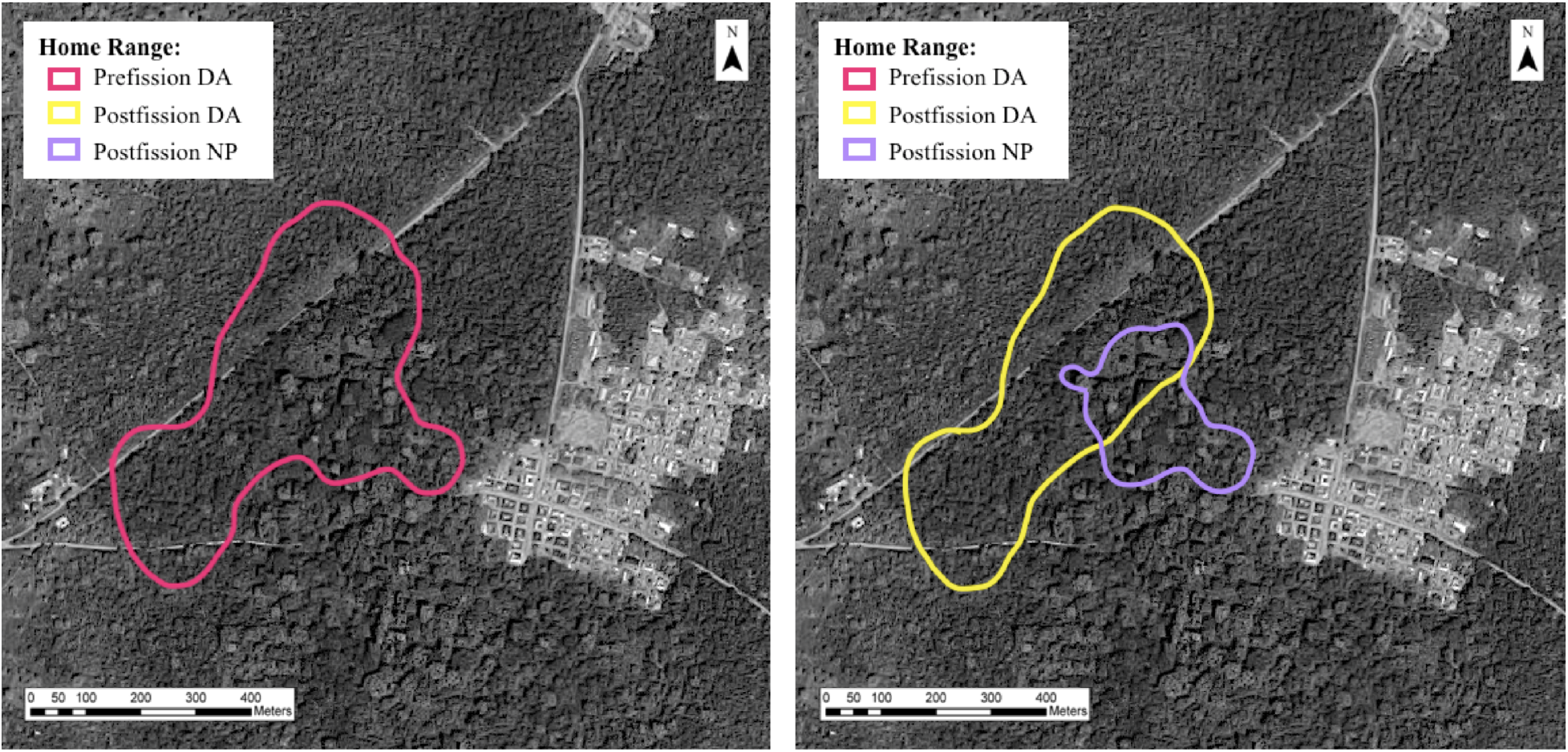
Home range distributions of the DA and NP groups prior to and after the fission event. (A) The original DA group maintained a large home range in the summer of 2006. (B) The two product groups split the original home range by the summer of 2007.

The original group was broken down into two groups for the sake of analysis—those female individuals which eventually split into the DA group and those females that split into the NP group. Average Bray-Curtis dissimilarity was 0.527 (SD +/- 0.046) for the two groups before the fission event, while it was 0.538 (SD +/- 0.048) for the two social groups after the fission event. In the PERMANOVA, age class (p = 0.378, df = 1, F = 1.028), collection site (p = 0.482, df = 2, F = 0.992), and reproductive status (p = 0.790, df = 2, F = 0.860) were not significant predictors of Bray-Curtis dissimilarity, but social group was a significant predictor (p = 0.001, df = 3, F = 4.416). Pairwise comparisons indicated that Bray-Curtis dissimilarity between the two social groups was not significant prior to the fission (p = 0.085, n = 12, pseudo F = 1.23), while differences after the fission were significant (p = 0.02, n = 12, pseudo F = 1.62). Significant differences were also found between the 2006 and 2007 sampling periods for all groups sampled (p < 0.01, n = 24, pseudo F = 2.19). Group membership explained 7.0% of the variation in gut microbiome diversity, while the largest predictor of variation was sampling year (2006 vs. 2007), explaining 31.4% of the variation in diversity (Fig. 2).

**Figure 2.**
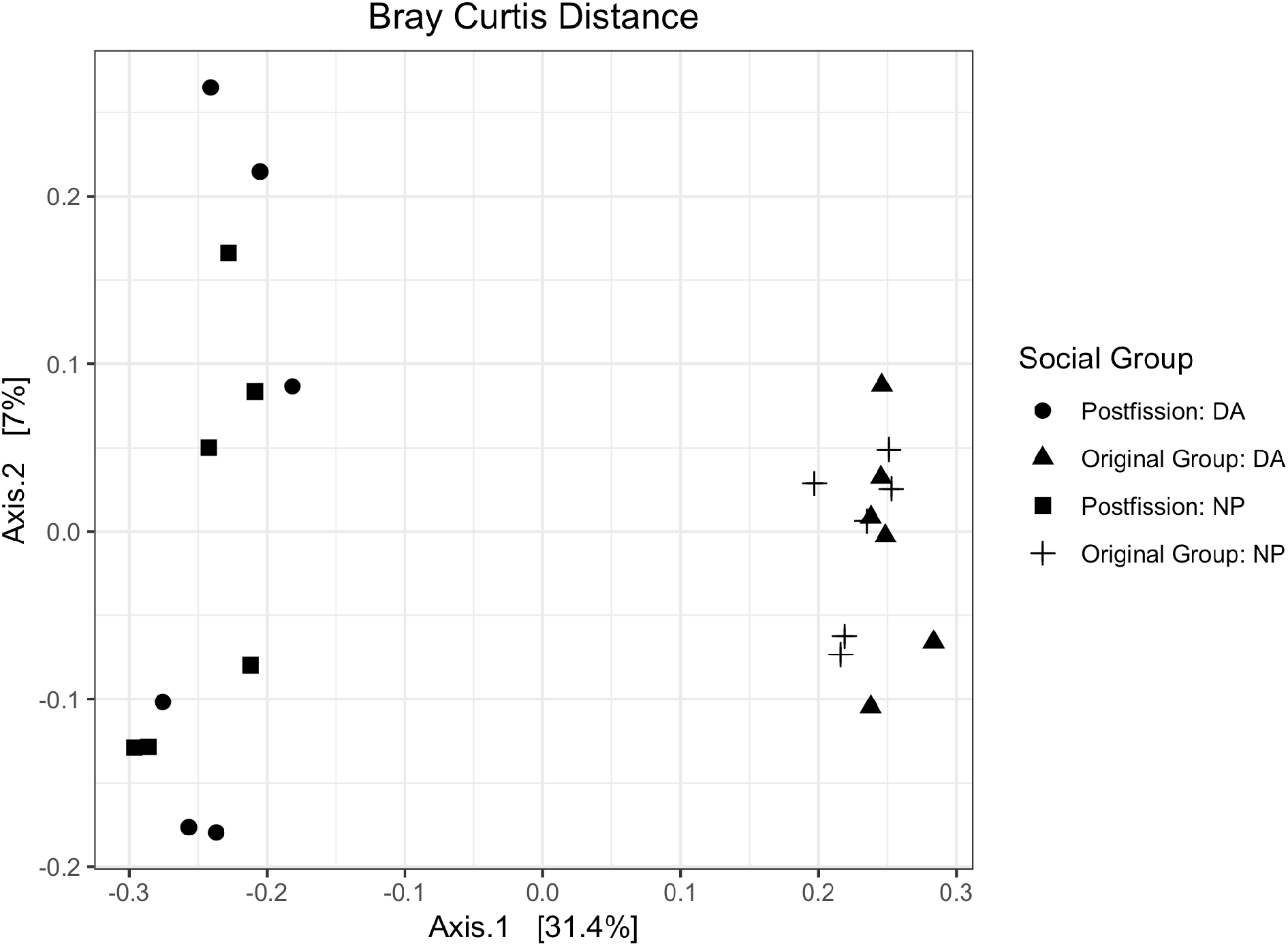
Post-fission group membership predicts Bray-Curtis dissimilarity of the NP and DA groups. No significant difference in gut microbial diversity was observed prior to the fission event (p = 0.085, n = 12, pseudo F = 1.23), while less than nine months after the fission event these groups showed unique microbial signatures (p = 0.02, n = 12, pseudo F = 1.62). Significant differences were found in gut microbial diversity between all groups across the years sampled (p < 0.01, n = 24, pseudo F = 2.19).

The average pairwise-relatedness among females in the social group prior to the fission was 0.153, while it was 0.232 in the post-fission NP group and 0.125 in the post-fission DA group (Fig. 3). This average pairwise-relatedness increased by 0.079 in the NP social group from the original DA group, suggesting that this fission event may have happened partially along lines of female relatedness, with a group of more closely-related female individuals splitting off to form the new NP group. However, the average pairwise-relatedness decreased by 0.028 between the pre-fission group and the new DA group. The pairwise-relatedness was not statistically different between any of the groups (p = 0.103, df = 2, χ^2^ = 4.5518). The Mantel test comparing Bray-Curtis dissimilarity and relatedness showed no statistically significant correlation between these variables (r < 0.01, p = 0.494).

**Figure 3.**
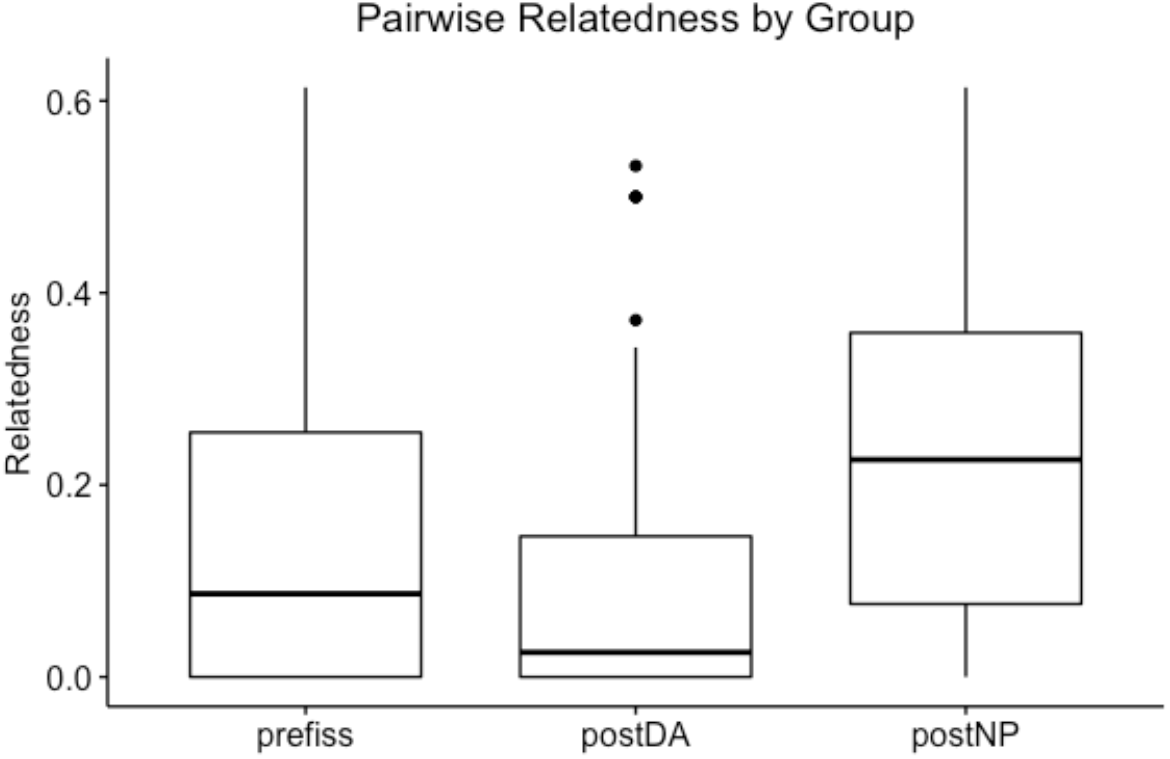
Pairwise-relatedness as calculated using 17 STR loci are represented by social group. On average, pairwise-relatedness increased by 0.079 in the NP social group from the original group. However, the pairwise-relatedness between groups was not statistically different (p=0.103, df=2, χ^2^=4.55).

On the whole, individual gut samples were vastly dominated by Firmicutes (57-78%) and Bacteriodes (2-13%). The next most prevalent phyla across samples were Tenericutes (3-10%) and Verucomicrobia (<1-16%). Linear discriminant effect analysis (LefSe) found no differences between the groups prior to the fission and three genera to differ between the two new social groups after the fission: *Parabacteroides, Coprococcus,* and *Porphyromonadaceae* (Fig. 4).

**Figure 4.**
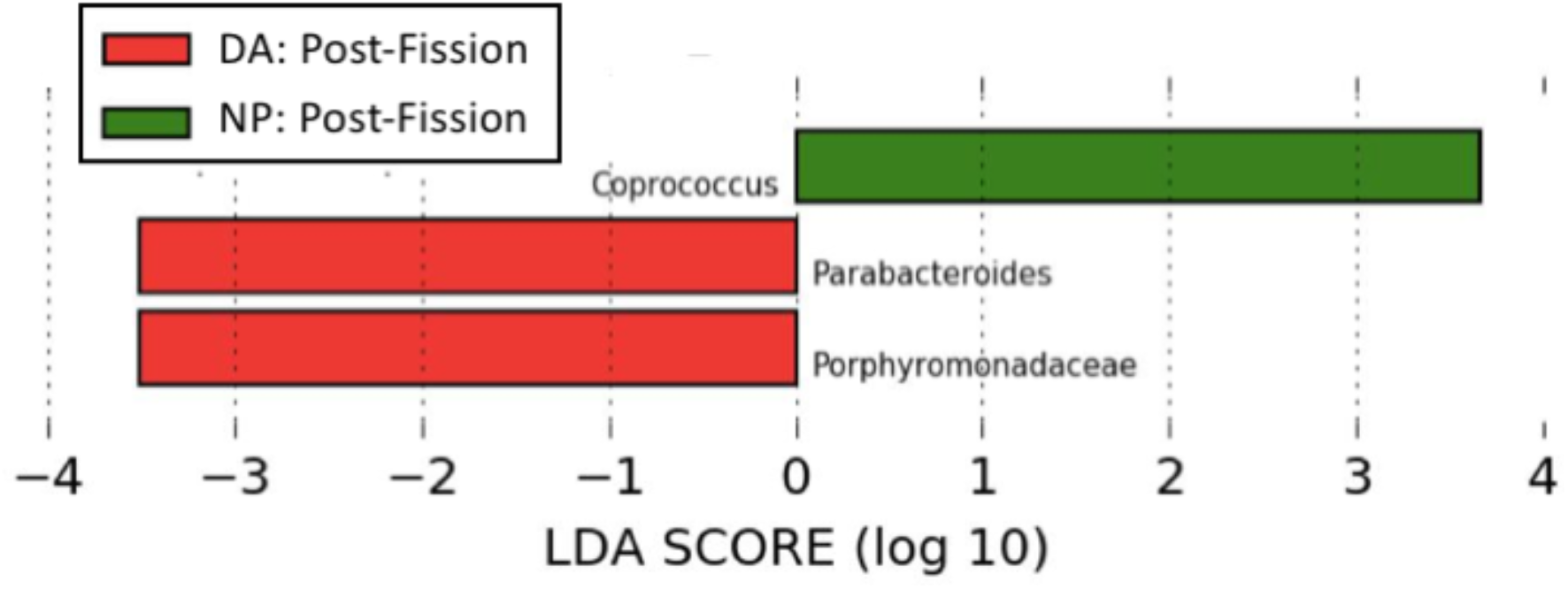
LDA scores for taxa differing significantly between product group. Three genera were found in different relative abundance between these two groups. Linear effect size analysis (LefSe) was run for the groups at a KW alpha value of 0.01 and an LDA score of 3.0.

## DISCUSSION

Distinct gut microbiota characterize different social groups across a wide range of taxa (Bennett *et al*., 2016; Degnan *et al.,* 2012; Tung *et al.,* 2015; Amato *et al.,* 2017). Examining the process and the time scale over which these divergences occur is important to understand the influence that social context can exert on gut microbiome assembly. Some recent research has focused on the time scale across which an individual’s microbiome converges with that of a new social group. Grieniesen *et al*. (2017) found that the longer an immigrant male baboon resided in a new social group, the more closely his core and non-core microbiomes resembled those of the adult members of that group, suggesting that the process of group convergence takes place over a span of months to years. Amaral *et al*. (2017) also found that, when newly-weaned infants joined new social groups, their gut microbiomes converged to resemble their new groups within two weeks. Our study focuses on the time scale across which social groups diverge from one another. We found distinct gut microbial signatures to characterize two daughter groups of colobus less than nine months after the fission event that resulted in these groups. Prior to this fission, the same individuals did not harbor distinct microbial communities, although the difference between them did approach significance. This trend may be due to sampling the original group at a timepoint during the initial stages of the fission event. Taken together, this finding both indicates that distinct gut microbial profiles can emerge in two new social groups in less than nine months and suggests that the process of group-specific microbial divergence may begin prior to the establishment of those groups. Additional timepoints leading up to and following a fission event are needed to more finely map the timescale across which these communities diverge and to better understand the mechanisms driving divergence.

Hosts can gain microbes through changes in social context, such as alterations in direct and indirect interactions with conspecifics that provide access to different microbes, potentially affecting gut microbiome composition (Lombardo *et al*., 2008). In social animals, changes in group composition, size, and social networks could all contribute to this type of shift. In this study, group composition of females before and after the fission remained similar overall (Table 1); thus the number and age structure of females in each fission product are unlikely to be driving our results. There were, however, other changes in social context in DA social group during the post-fission field season, including two males immigrating to DA group and one infant dying. Further investigation into how changes in social environment and social stress affect the gut microbiome are required to determine how these events may have influenced the observed gut microbiome divergence.

Other factors potentially contributing to the observed microbial shifts are diet and/or ranging patterns. While diet has been suggested as a primary driver in structuring the gut microbiome (Muegge *et al*., 2011; Amato *et al.,* 2014; Hale *et al*., 2018), diet has not been found to explain differences in gut microbial beta diversity between individuals and groups in our population (Wikberg *et al.,* 2017; Wikberg unpublished data). Alternatively, despite all females in NP and DA group using a distinct part of their group’s home range as well as a large overlap zone between the two groups (Fig. 1) and collection site being a non-significant predictor variable for gut microbial dissimilarity, we cannot rule out the effects of habitat use on the observed shifts. While product groups ranged in subsets of the original range that included both woodland and mature forest, DA ranged primarily in woodland areas after the fission while NP’s new range tended toward the mature forest. Because even small changes in environment can expose animals to new reservoirs of environmentally-derived microbes, it is possible that the divergence observed between the two product groups is in part driven by spatial distribution and habitat use.

As has been reported in other folivorous species, individual gut samples were vastly dominated by Firmicutes and Bacteriodes with low but consistent proportions of Tenericutes and Verucomicrobia (Yildirim *et al*., 2010; Amato *et al*., 2016). Linear discriminant analysis (LefSe) revealed three genera to differ between the two product groups of this study. The new DA group was found to have relatively more *Porphyromonadaceae* and *Parabacteriodes* than the NP group after the fission, while the genus *Coprococcus* was found to be at greater prevalence in the NP group than the new DA group. The *Coprococcus* genus is in the order Clostridiales, which can assist in the degradation of plant material and is likely to reflect the folivorous diet of these animals (Barelli *et al*., 2015). It has been found at differential abundances in different social groups of baboons, suggesting that this might be a genus with a strong propensity for social transmission (Grieneisen *et al*., 2017). This genus is also commonly used to gauge individual gut health, and decreased levels of *Coprococcus* have been shown to accompany a stress response (Derrien *et al*., 2015), which could be related to the changes in social context and/or increased ranging in woodland habitat seen in the DA social group. Because quadrats characterized as “woodland” at BFMS have been previously found to have fewer large trees, less species diversity, and a lower basal area of colobus food trees than those in the interior of the forest (Teichroeb & Sicotte, 2018), it is possible that the DA group’s increased ranging in this type of habitat may partially account for the elevated levels of *Coprococcus* observed in this group. While our analyses showed that collection site (woodland forest vs. mature forest) was not found to be a significant predictor of variation in this study, more detailed study on the effects of ranging patterns and habitat use are required.

This fission event resulted in an increase in average pairwise-relatedness for the NP and a decrease for the DA group, although there were no significant differences in mean relatedness between the original group and the post-fission groups. This is a common phenomenon in animals that disperse by group fission, and an increase in relatedness in fission product groups has been demonstrated widely across primate species (Widdig *et al*., 2006; Snyder-Mackler *et al*., 2014). Although genetic variation can play a significant role in shaping the diversity of the gut microbiome (Goodrich *et al*., 2014), previous studies have found little evidence for a strong role of host genetics in structuring the microbial communities of wild primates (Degnan *et al*., 2012; Amato *et al*., 2017; Spor *et al*., 2011). In this particular data set, no correlation existed between beta diversity and relatedness. Taken together, our results suggest that even though NP group contained some close female kin dyads, relatedness did not play a significant role in structuring the differences in beta diversity seen between the two groups.

Finally, the largest proportion of variation between groups in this study was explained by year, rather than group membership. Previous studies have found temporal variation in the gut microbiomes of other folivorous primates to change in response to seasonal changes in food availability (Amato *et al*., 2015; Springer *et al*. 2017), which is consistent with past observations in this study population (Wikberg *et al*., 2016, Wikberg unpublished data). However, because longitudinally collected samples in this study were all from the wet season, the observed differences would need to be explained by some aspect of interannual variation in food availability during the same season. Further work is needed to clarify this possibility, including more sampling points between years and seasons as well as detailed data on changes in diet and food availability through time.

Overall, we used a longitudinal approach that provides a new perspective into how social groups acquire distinct gut microbial communities and the time period over which these divergent communities establish. This has significant consequences for understanding the role of social context in shaping the unique microbial signatures associated with distinct social groups across a wide variety of taxa. Further work is recommended into more finely mapping the timescales and factors that result in this divergence, especially within the context of the potentially adaptive effects of this recurrent, social-context dependent trend.

## ACKNOWLEDGEMENTS

We confirm that all research protocols reported in this manuscript were reviewed and approved by an appropriate institution and/or governmental agency that regulates research with animals, all research reported in this manuscript adhered to the legal requirements of the country in which the work took place, and that the research adhered to the American Society of Primatologists (ASP) Principles for the Ethical Treatment of Non Human Primates. Specifically, we gained permission from the Ghana Wildlife Division, the management committee at BFMS, and the University of Calgary’s Animal Care Committee to conduct this study. Funding was granted by Alberta Ingenuity, American Society of Primatologists, International Primatological Society, Leakey Foundation, Natural Sciences and Engineering Research Council of Canada, Province of Alberta, Sweden-America Foundation, Wenner–Gren Foundation (8172), the University of Calgary, the University of Oregon’s O’Day Fellowship Program in Biological Sciences and Office of the Vice President for Research and Innovation, and the National Institute of General Medical Sciences (P50GM098911) via the META Center for Systems Biology. We thank two anonymous reviewers for their helpful feedback.

## REFERENCES

Amaral, W. Z., Lubach, G. R., Proctor, A., Lyte, M., Phillips, G. J., & Coe, C. L. (2017). Social influences on prevotella and the gut microbiome of young monkeys. Psychosomatic medicine, 79(8), 888–897. https://doi.org/10.1097/psy.0000000000000454

Amato, K. (2013). Co-evolution in context: The importance of studying gut microbiomes in wild animals. Microbiome Sci Med, 1(1), 10–29. https://doi.org/10.2478/micsm-2013-0002

Amato, K. R., Leigh, S. R., Kent, A., Mackie, R. I., Yeoman, C. J., Stumpf, R. M., … Garber, P. A. (2014). The Gut Microbiota Appears to Compensate for Seasonal Diet Variation in the Wild Black Howler Monkey (Alouatta pigra). Microbial Ecology, 69(2), 434–443. https://doi.org/10.1007/s00248-014-0554-7

Amato, K. R., Yeoman, C. J., Cerda, G., Schmitt, C. A., Cramer, J. D., Miller, M. E. B., … Leigh, S. R. (2015). Variable responses of human and non-human primate gut microbiomes to a Western diet. Microbiome, 3, 53. https://doi.org/10.1186/s40168-015-0120-7

Amato, K. R., Martinez-Mota, R., Righini, N., Raguet-Schofield, M., Corcione, F. P., Marini, E., … & Williams, L. (2016). Phylogenetic and ecological factors impact the gut microbiota of two Neotropical primate species. Oecologia, 180(3), 717–733. https://doi.org/10.1007/s00442-015-3507-z

Amato, K. R., Van Belle, S., Di Fiore, A., Estrada, A., Stumpf, R., White, B., … & Leigh, S. R. (2017). Patterns in Gut Microbiota Similarity Associated with Degree of Sociality among Sex Classes of a Neotropical Primate. Microbial Ecology, 74(1), 250–258. https://doi.org/10.1007/s00248-017-0938-6

Archie, E. A., & Tung, J. (2015). Social behavior and the microbiome. Current Opinion in Behavioral Sciences, 6, 28–34. https://doi.org/10.1016/j.cobeha.2015.07.008

Barbáchano, A., Fernández-Barral, A., Ferrer-Mayorga, G., Costales-Carrera, A., Larriba, M. J., & Muñoz, A. (2017). The endocrine vitamin D system in the gut. Molecular and cellular endocrinology, 453, 79–87. https://doi.org/10.1016/j.mce.2016.11.028

Bădescu, I., Sicotte, P., Ting, N., & Wikberg, E. C. (2015). Female parity, maternal kinship, infant age and sex influence natal attraction and infant handling in a wild colobine (Colobus vellerosus). American journal of primatology, 77(4), 376–387. https://doi.org/10.1002/ajp.22353

Barelli, C., Albanese, D., Donati, C., Pindo, M., Dallago, C., Rovero, F., … De Filippo, C. (2015). Habitat fragmentation is associated to gut microbiota diversity of an endangered primate: Implications for conservation. Scientific Reports, 5, 1–12. https://doi.org/10.1038/srep14862

Bennett, G., Malone, M., Sauther, M. L., Cuozzo, F. P., White, B., Nelson, K. E., … & Amato, K. R. (2016). Host age, social group, and habitat type influence the gut microbiota of wild ring-tailed lemurs (Lemur catta). American journal of primatology, 78(8), 883–892. https://doi.org/10.1002/ajp.22555

Bolger, A. M., Lohse, M., & Usadel, B. (2014). Trimmomatic: a flexible trimmer for Illumina sequence data. Bioinformatics, 30(15), 2114–2120. https://doi.org/10.1093/bioinformatics/btu170

Bonte, D., Van Dyck, H., Bullock, J. M., Coulon, A., Delgado, M., Gibbs, M., … & Schtickzelle, N. (2012). Costs of dispersal. Biological Reviews, 87(2), 290–312. https://doi.org/10.1111/j.1469-185X.2011.00201.x

Cho, I., & Blaser, M. J. (2012). The human microbiome: at the interface of health and disease. Nature Reviews Genetics, 13(4), 260. https://doi.org/10.1038/nrg3182

Costello, E. K., Stagaman, K., Dethlefsen, L., Bohannan, B. J., & Relman, D. A. (2012). The application of ecological theory toward an understanding of the human microbiome. Science, 336(6086), 1255–1262. https://doi.org/10.1126/science.1224203

Degnan, P. H., Pusey, A. E., Lonsdorf, E. V, Goodall, J., Wroblewski, E. E., & Wilson, M. L. (2012). Factors associated with the diversification of the gut microbial communities within chimpanzees from Gombe National Park. Proceedings of the National Academy of Sciences (USA), 109, 13034–13039. https://doi.org/10.1073/pnas.1110994109

Derrien, M., Johan, E.T., & Vileg, H. (2015). Fate, activity, and impact of ingested bacteria within the human gut microbiota. Trends in Microbiology, 23(6), 354–66. https://doi.org/10.1016/j.tim.2015.03.002

Dunbar, R. I. M., MacCarron, P., & Robertson, C. (2018). Trade-off between fertility and predation risk drives a geometric sequence in the pattern of group sizes in baboons. Biology letters, 14(3), 20170700. https://doi.org/10.1098/rsbl.2017.0700

Ezenwa, V. O., Gerardo, N. M., Inouye, D. W., Medina, M., & Xavier, J. B. (2012). Animal behavior and the microbiome. Science, 338(6104), 198–199. https://doi.org/10.1126/science.1227412

Goodrich, J. K., Waters, J. L., Poole, A. C., Sutter, J. L., Koren, O., Blekhman, R., … & Spector, T. D. (2014). Human genetics shape the gut microbiome. Cell, 159(4), 789–799. https://doi.org/10.1016/j.cell.2014.09.053

Grieneisen, L. E., Livermore, J., Alberts, S., Tung, J., & Archie, E. A. (2017). Group Living and Male Dispersal Predict the Core Gut Microbiome in Wild Baboons. Integrative and Comparative Biology, 0(0), 1–16. https://doi.org/10.1093/icb/icx046

Hale, V. L., Tan, C. L., Niu, K., Yang, Y., Knight, R., Zhang, Q., … & Amato, K. R. (2018). Diet versus phylogeny: a comparison of gut microbiota in captive Colobine monkey species. Microbial ecology, 75(2), 515–527. https://doi.org/10.1007/s00248-017-1041-8

Hall, J. B., & Swaine, M. D. (2013). Distribution and ecology of vascular plants in a tropical rain forest: forest vegetation in Ghana (Vol. 1). Springer Science & Business Media.

Henzi, S. P., Lycett, J. E., & Weingrill, T. (1997). Cohort size and the allocation of social effort by female mountain baboons. Animal Behaviour, 54, 1235–1243. https://doi.org/10.1006/anbe.1997.0520

Huttenhower, C., Gevers, D., Knight, R., Abubucker, S., Badger, J. H., Chinwalla, A. T., … & White, O. (2012). Structure, function and diversity of the healthy human microbiome. Nature, 486(7402), 207–214. https://doi.org/10.1038/nature11234

Kankam, B. O., & Sicotte, P. (2013). The effect of forest fragment characteristics on abundance of Colobus vellerosus in the Forest-Savanna Transition Zone of Ghana. Folia Primatologica, 84(2), 74–86. https://doi.org/10.1159/000348307

Leamy, L. J., Kelly, S. A., Nietfeldt, J., Legge, R. M., Ma, F., Hua, K., … & Pomp, D. (2014). Host genetics and diet, but not immunoglobulin A expression, converge to shape compositional features of the gut microbiome in an advanced intercross population of mice. Genome biology, 15(12), 552. https://doi.org/10.1186/s13059-014-0552-6

Lombardo, M. P. (2008). Access to mutualistic endosymbiotic microbes: an underappreciated benefit of group living. Behavioral Ecology and Sociobiology, 62(4), 479–497. https://doi.org/10.1007/s00265-007-0428-9

Markham, A. C., Gesquiere, L. R., Alberts, S. C., & Altmann, J. (2015). Optimal group size in a highly social mammal. Proceedings of the National Academy of Sciences, 112(48), 14882–14887. https://doi.org/10.1073/pnas.1517794112

McCord, A. I., Chapman, C. A., Weny, G., Tumukunde, A., Hyeroba, D., Klotz, K., … & Leigh, S. R. (2014). Fecal microbiomes of non-human primates in Western Uganda reveal species-specific communities largely resistant to habitat perturbation. American journal of primatology, 76(4), 347–354. https://doi.org/10.1002/ajp.22238

McMurdie, P. J., & Holmes, S. (2013). phyloseq: an R package for reproducible interactive analysis and graphics of microbiome census data. PloS one, 8(4), e61217. https://doi.org/10.1371/journal.pone.0061217

Moeller, A. H., Foerster, S., Wilson, M. L., Pusey, A. E., Hahn, B. H., & Ochman, H. (2016). Social behavior shapes the chimpanzee pan-microbiome. Science Advances, 2(1), e1500997. https://doi.org/10.1126/sciadv.1500997

Morin, P. A., Chambers, K. E., Boesch, C., & Vigilant, L. (2001). Quantitative polymerase chain reaction analysis of DNA from noninvasive samples for accurate microsatellite genotyping of wild chimpanzees (Pan troglodytes verus). Molecular ecology, 10(7), 1835–1844. https://doi.org/10.1046/j.0962-1083.2001.01308.x

Muegge, B. D., Kuczynski, J., Knights, D., Clemente, J. C., González, A., Fontana, L., … & Gordon, J. I. (2011). Diet drives convergence in gut microbiome functions across mammalian phylogeny and within humans. Science, 332(6032), 970–974. https://doi.org/10.1126/science.1198719

Perofsky, A. C., Lewis, R. J., Abondano, L. A., Di Fiore, A., & Meyers, L. A. (2017). Hierarchical social networks shape gut microbial composition in wild Verreaux’s sifaka. Proc. R. Soc. B, 284(1868), 20172274. https://doi.org/10.1098/rspb.2017.2274

Rollins, L.A., Browning, L.E., Holleley, C.E., Savage, J.L., Russell, A.F., Griffith, S.C. (2012). Building genetic networks using relatedness information: a novel approach for the estimation of dispersal and characterization of group structure in social animals. Mol. Ecol. 21, 1727–1740. https://doi.org/10.1111/j.1365-294X.2012.05492.x

Saj, T. L., Teichroeb, J. A., & Sicotte, P. (2005). The population status of the ursine colobus (Colobus vellerosus) at Boabeng-Fiema, Ghana. Commensalism and conflict: the human primate interface (Paterson, JD & Wallis, J., eds). American Society of Primatologists, Norman, OK, 350–375.

Saj, T. L., & Sicotte, P. (2007). Scramble competition among Colobus vellerosus at Boabeng-Fiema, Ghana. International Journal of Primatology, 28(2), 337–355. https://doi.org/10.1007/s10764-007-9125-9

Saj, T. L., & Sicotte, P. (2007). Predicting the competitive regime of female Colobus vellerosus from the distribution of food resources. International Journal of Primatology, 28(2), 315–336. https://doi.org/10.1007/s10764-007-9124-x

Segata, N., Izard, J., Waldron, L., Gevers, D., Miropolsky, L., Garrett, W. S., & Huttenhower, C. (2011). Metagenomic biomarker discovery and explanation. Genome biology, 12(6), R60. https://doi.org/10.1186/gb-2011-12-6-r60

Sicotte, P., & Macintosh, A. J. (2004). Inter-group encounters and male incursions in Colobus vellerosus in central Ghana. Behaviour, 141(5), 533–553. https://doi.org/10.1163/1568539041166717

Sicotte, P., Teichroeb, J. A., Vayro, J. V., Fox, S. A., Bădescu, I., & Wikberg, E. C. (2017). The influence of male takeovers on female dispersal in Colobus vellerosus. American journal of primatology, 79(7), e22436. https://doi.org/10.1002/ajp.22436

Snyder-Mackler, N., Alberts, S. C., & Bergman, T. J. (2014). The socio-genetics of a complex society: female gelada relatedness patterns mirror association patterns in a multilevel society. Molecular ecology, 23(24), 6179–6191. https://doi.org/10.1111/mec.12987

Spor, A., Koren, O., & Ley, R. (2011). Unravelling the effects of the environment and host genotype on the gut microbiome. Nature Reviews Microbiology, 9(4), 279. https://doi.org/10.1038/nrmicro2540

Springer, A., Fichtel, C., Al-Ghalith, G. A., Koch, F., Amato, K. R., Clayton, J. B., … & Kappeler, P. M. (2017). Patterns of seasonality and group membership characterize the gut microbiota in a longitudinal study of wild Verreaux’s sifakas (Propithecus verreauxi). Ecology and evolution, 7(15), 5732–5745. https://doi.org/10.1002/ece3.3148

Strum, S. C. (2012). Darwin’s monkey: Why baboons can’t become human. American journal of physical anthropology, 149(S55), 3–23. https://doi.org/10.1002/ajpa.22158

Sueur, C., & Maire, A. (2014). Modelling animal group fission using social network dynamics. PloS one, 9(5), e97813. https://doi.org/10.1371/journal.pone.0097813

Teichroeb, J. A., Wikberg, E. C., & Sicotte, P. (2009). Female dispersal patterns in six groups of ursine colobus (*Colobus vellerosus*): infanticide avoidance is important. Behaviour, 146(4), 551–582. https://doi.org/10.1163/156853909X426363

Teichroeb, J. A., Wikberg, E. C., & Sicotte, P. (2011). Dispersal in male ursine colobus monkeys (Colobus vellerosus): influence of age, rank and contact with other groups on dispersal decisions. Behaviour, 148(7), 765–793. https://doi.org/10.1163/000579511X577157

Teichroeb, J. A., & Sicotte, P. (2012). Cost-free vigilance during feeding in folivorous primates? Examining the effect of predation risk, scramble competition, and infanticide threat on vigilance in ursine colobus monkeys (Colobus vellerosus). Behavioral Ecology and Sociobiology, 66(3), 453–466. https://doi.org/10.1007/s00265-011-1292-1

Teichroeb, J. A., & Sicotte, P. (2018). Cascading competition: the seasonal strength of scramble influences between-group contest in a folivorous primate. Behavioral Ecology and Sociobiology, 72(1), 6. https://doi.org/10.1007/s00265-017-2418-x

Tung, J., Barreiro, L. B., Burns, M. B., Grenier, J. C., Lynch, J., Grieneisen, L. E., … & Archie, E. A. (2015). Social networks predict gut microbiome composition in wild baboons. Elife, 4, e05224. https://doi.org/10.7554/eLife.05224.002

Turnbaugh, P. J., Hamady, M., Yatsunenko, T., Cantarel, B. L., Duncan, A., Ley, R. E., … & Egholm, M. (2009). A core gut microbiome in obese and lean twins. Nature, 457(7228), 480. https://doi.org/10.1038/nature07540

Wang, J. (2011). COANCESTRY: A program for simulating, estimating and analysing relatedness and inbreeding coefficients. Molecular Ecology Resources, 11, 141–145. https://doi.org/10.1111/j.1755-0998.2010.02885.x

Widdig, A., Nürnberg, P., Bercovitch, F. B., Trefilov, A., Berard, J. B., Kessler, M. J., … & Krawczak, M. (2006). Consequences of group fission for the patterns of relatedness among rhesus macaques. Molecular Ecology, 15(12), 3825–3832. https://doi.org/10.1111/j.1365-294X.2006.03039.x

Wikberg, E. C., Sicotte, P., Campos, F. A., & Ting, N. (2012). Between-Group Variation in Female Dispersal, Kin Composition of Groups, and Proximity Patterns in a Black-and-White Colobus Monkey (Colobus vellerosus). PLoS ONE, 7(11), 1–14. https://doi.org/10.1371/journal.pone.0048740

Wikberg, E. C., Teichroeb, J. A., Bădescu, I., & Sicotte, P. (2013). Individualistic female dominance hierarchies with varying strength in a highly folivorous population of black-and-white colobus. Behaviour, 150(3-4), 295–320. https://doi.org/10.1163/1568539X-00003050

Wikberg, E. C., Ting, N., & Sicotte, P. (2014). Familiarity is more important than phenotypic similarity in shaping social relationships in a facultative female dispersed primate, Colobus vellerosus. Behavioural processes, 106, 27–35. https://doi.org/10.1016/j.beproc.2014.04.002

Wikberg, E. C., Ting, N., & Sicotte, P. (2014). Kinship and similarity in residency status structure female social networks in black-and-white colobus monkeys (*Colobus vellerosus*). American Journal of Physical Anthropology, 153(3), 365–376. https://doi.org/10.1002/ajpa.22435

Wikberg, E. C., Ting, N., & Sicotte, P. (2015). Demographic Factors Are Associated with Intergroup Variation in the Grooming Networks of Female Colobus (Colobus vellerosus). International Journal of Primatology, 36(1), 124–142. https://doi.org/10.1007/s10764-015-9816-6

Wikberg, E.C., Christie, D.M., Sicotte, P., & Ting, N. (2017, August 21-27). Between and within group variation in the gut microbiome of a black-and-white colobus monkey (*Colobus vellerosus*). Paper presented at the International Primatologcal Society Congress.

Wikberg, E. C., Christie, D.M., Campos, F. A., Sicotte, P., & Ting, N. (2017). The link between social networks and gut microbial composition in black-and-white colobus (Colobus vellerosus). In American Journal of Physical Anthropology, 162(S64), 409–409.

Williams, B.L., Hornig, M., Parekh, T., Lipkin, W.I. (2012). Application of novel PCR-based methods for detection, quantitation, and phylogenetic characterization of Sutterella species in intestinal biopsy samples from children with autism and gastro-intestinal disturbances. mBio 3, e00261–11. https://doi.org/10.1128/mBio.00261-11

Wong, S. N. P., & Sicotte, P. (2006). Population size and density of Colobus vellerosus at the Boabeng-Fiema Monkey Sanctuary and surrounding forest fragments in Ghana. American Journal of Primatology, 68, 465–476. https://doi.org/10.1002/ajp.20242

Wu, G. D., Chen, J., Hoffmann, C., Bittinger, K., Chen, Y. Y., Keilbaugh, S. A., … & Sinha, R. (2011). Linking long-term dietary patterns with gut microbial enterotypes. Science, 334(6052), 105–108. https://doi.org/10.1126/science.1208344

Yildirim, S., Yeoman, C. J., Sipos, M., Torralba, M., Wilson, B. A., Goldberg, T. L., … & Nelson, K. E. (2010). Characterization of the fecal microbiome from non-human wild primates reveals species specific microbial communities. PloS one, 5(11), e13963. https://doi.org/10.1371/journal.pone.0013963

